# Structural and functional analysis of female sex hormones against SARS-Cov2 cell entry

**DOI:** 10.1101/2020.07.29.227249

**Authors:** Jorge Alberto Aguilar-Pineda, Mazen Albaghdadi, Wanlin Jiang, Karin J. Vera Lopez, Gonzalo Davila Del-Carpio, Badhin Gómez Valdez, Mark E. Lindsay, Rajeev Malhotra, Christian L. Lino Cardenas

**Affiliations:** Cardiovascular Research Center, Cardiology Division, Massachusetts General Hospital, Harvard Medical School, Boston, Massachusetts, USA; Laboratory of Genomics and Neurovascular Diseases, Vicerrectorado de Investigación, Universidad Católica de Santa María, Arequipa, Peru

## Abstract

Emerging evidence suggests that males are more susceptible to severe infection by the SARS-CoV-2 virus than females. A variety of mechanisms may underlie the observed gender-related disparities including differences in sex hormones. However, the precise mechanisms by which female sex hormones may provide protection against SARS-CoV-2 infectivity remains unknown. Here we report new insights into the molecular basis of the interactions between the SARS-CoV-2 spike (S) protein and the human ACE2 receptor. We further observed that glycosylation of the ACE2 receptor enhances SARS-CoV-2 infectivity. Importantly estrogens can disrupt glycan-glycan interactions and glycan-protein interactions between the human ACE2 and the SARS-CoV2 thereby blocking its entry into cells. In a mouse model, estrogens reduced ACE2 glycosylation and thereby alveolar uptake of the SARS-CoV-2 spike protein. These results shed light on a putative mechanism whereby female sex hormones may provide protection from developing severe infection and could inform the development of future therapies against COVID-19.

## Introduction

The novel coronavirus disease 2019 (COVID-19) global pandemic caused by infection with the severe acute respiratory syndrome coronavirus 2 (SARS-Cov2) virus has infected nearly 15 million people worldwide resulting in nearly 600,000 deaths as July 21, 2020^1^. Emerging data suggests that males are more susceptible to COVID-19 infection and are at higher risk of critical illness and death than females^2–4^. There has been consistent evidence of an increased case fatality rate (CFR) among males in nearly every country with available sex-disaggregated data including Peru, France, Greece, Italy, Mexico, Pakistan, Philippines and Spain amounting to a 1.7 times higher CFR than females^5^.

Understanding the mechanisms underlying enhanced COVID-19 susceptibility and disease severity in males is key to developing new therapies and guiding vaccine development. Changes in sex hormone concentration over an individual’s lifetime and associated risk of comorbid conditions, such as cardiovascular diseases, may also contribute to variability in disease susceptibility and severity^6^. It has been postulated that the male-biased sex divergence in COVID-19 deaths could be, in part, explained by the strict relationship between sex hormones and the expression of the entry receptor for SARS-CoV2, the angiotensin converting enzyme 2 (ACE2) receptor^3,7^. Molecular studies have demonstrated that the male hormone testosterone regulates the expression of ACE2 and the transmembrane serine protease 2 (TMPRSS2) which is an androgen-responsive serine protease that cleaves the SARS-CoV-2 spike (S) protein and facilitates viral entry via ACE2 binding^8–10^. Androgen-driven upregulation of ACE2 levels may therefore be associated with increased vulnerability to severe infections in male patients with COVID-19. Paradoxically, ACE2 plays an important role in lung protection during injury which is attenuated by the binding of SARS-CoV-2^11^.

The presence of a male-biased dependence in COVID-19 susceptibility may imply the presence of a protective factor against SARS-CoV-2 infectivity in women. In addition to the ability of sex hormones to modulate expression of proteins related to entry into host cells, both estrogens and androgens are also able to directly modulate immune cell function via receptor-mediated effects^12,13^. Additionally, sex chromosomes may mediate more favorable outcomes among women compared to men affected with COVID-19^14^. X-linked genes associated with immune function tend to be expressed more often in females who generally have two X chromosomes compared to males^15^.

Additional clues to the possible protective effects of estrogens have been suggested by differences in dietary patterns among countries with different CFRs^16^. Interestingly, countries with the lowest CFRs including Japan and Korea are the largest consumers of isoflavones-based foods, also known as phytoestrogens, that may also mediate favorable effects on ACE2 expression and therefore COVID-19 risk^17–19^. The observation that females and those individuals consuming higher levels of isoflavones may be protected from COVID-19 infection and adverse consequences indicates a potential protective role of estrogens against SARS-CoV-2.

Here, we examine the role of two estrogen molecules (17β-diol and S-equol) to modulate the ACE2-dependent membrane fusion protein and reduce cell entry of the SARS-CoV2 spike protein into lung cells. To the best of our knowledge, we report new findings regarding the importance of molecular interactions between hACE2 and the viral spike (S) protein. Furthermore, we provide insights into the molecular basis for our observations that estrogens impair SARS-CoV2 entry and highlight the potential for estrogens as an agent in patients with COVID-19.

## Results

### Glycosylation site-mapping of human ACE2 and SARS-CoV-2 spike interactions

Recent studies^20,21^ have shown the ability of the SARS-CoV2 virus to utilize a highly glycosylated spike (S) protein to elude the host’s immune system and bind to its target membrane receptor, ACE2, thus enabling entry into human cells. Based on the structural complementarity and steric impediments between the S protein and human ACE2 (hACE2) protein membranes, we mapped the glycosylation sites of both models^21–24^ and performed molecular dynamics simulations (MDS) by 250 ns to stabilize the glycosylated SARS-CoV2 spike (S) and hACE2 complex (suppl. Table 1., suppl. Figure 1 and Figure 1a). These analyses revealed that glycosylation of the ACE2 protein increases the affinity of the virus S protein to interact with the receptor via glycan-glycan interactions, glycan-protein interactions, hydrogen, and hydrophobic bonds (suppl. Table 2. and Figure 1b). Notably, glycan-glycan interactions occur between the ACE2 glycan at N322 and N546 and glycans found on the spike’s receptor binding domain (S-RBD) at N165 and N343 (Figure 1c, left panel). Despite the close interaction between ACE2 and S-RBD glycans, their affinity to anchor with highly negatively charged molecules such as the ACE2 protein remains unalterable (Figure 1c right panel) suggesting that glycan and electrostatic-dependent surface tethering may represent a plausible mechanism for ACE-S-RBD binding and cell infection. The glycan-protein interactions occur between the ACE2 glycan at N53 and the residues of the S-RBD at N437, S438, N439, L441, V445, G446, V483, Q498, T500 and Q506 (Figure 1d). While ACE2 residues at D38, Y41, W48 and G326 form hydrogen bonds with residues of the S-RBD at N440, D442, S443, N450 and E484 (Figure 1f). Multiple distinct clusters of hydrophobic residues at the ACE2 surface were also found to interact with the S-RBD protein (suppl. Figure 2). Importantly one key hydrophobic region on the ACE2 surface at T334 interacts with five residues of the S-RBD (P479, G485, F486, G488 and Y489), (Figure 1g). Given the insights afforded by our in silico MDS experiments, we sought to explore the impact of ACE2 glycosylation on S-RBD cell entry using cultured human umbilical vein endothelial cells (HUVECs). A variety of saccharide substrates were utilized for their ability to modulate glycosylation profiles in cells. The glycosylation pattern of the endogenous ACE2 was increased in nearly all treated cells (Figure 1h). Notably co-incubation of HUVECs with 10ug of recombinant S-RBD (rS-RBD) protein revealed that glucose (25mM) pre-treatment was associated with the greatest degree of rS-RBD entry into the cells by ~8 fold compared with hypoglycemic media (HBSS or Optimen) cells (Figure 1g). This model indicates that glycosylated residues surrounding the cavity at the top of the ACE2 molecule could increase binding by the S-RBD. Given the possibility that occupancy at glycosylated residues or S-RBD binding sites by estrogens could modify the affinity of the SARS-CoV2 virus and alter entry into the cell thereby reducing infectivity, we sought to further examine these interactions using a range of complementary experimental approaches (see Table S1).

**Figure 1.**
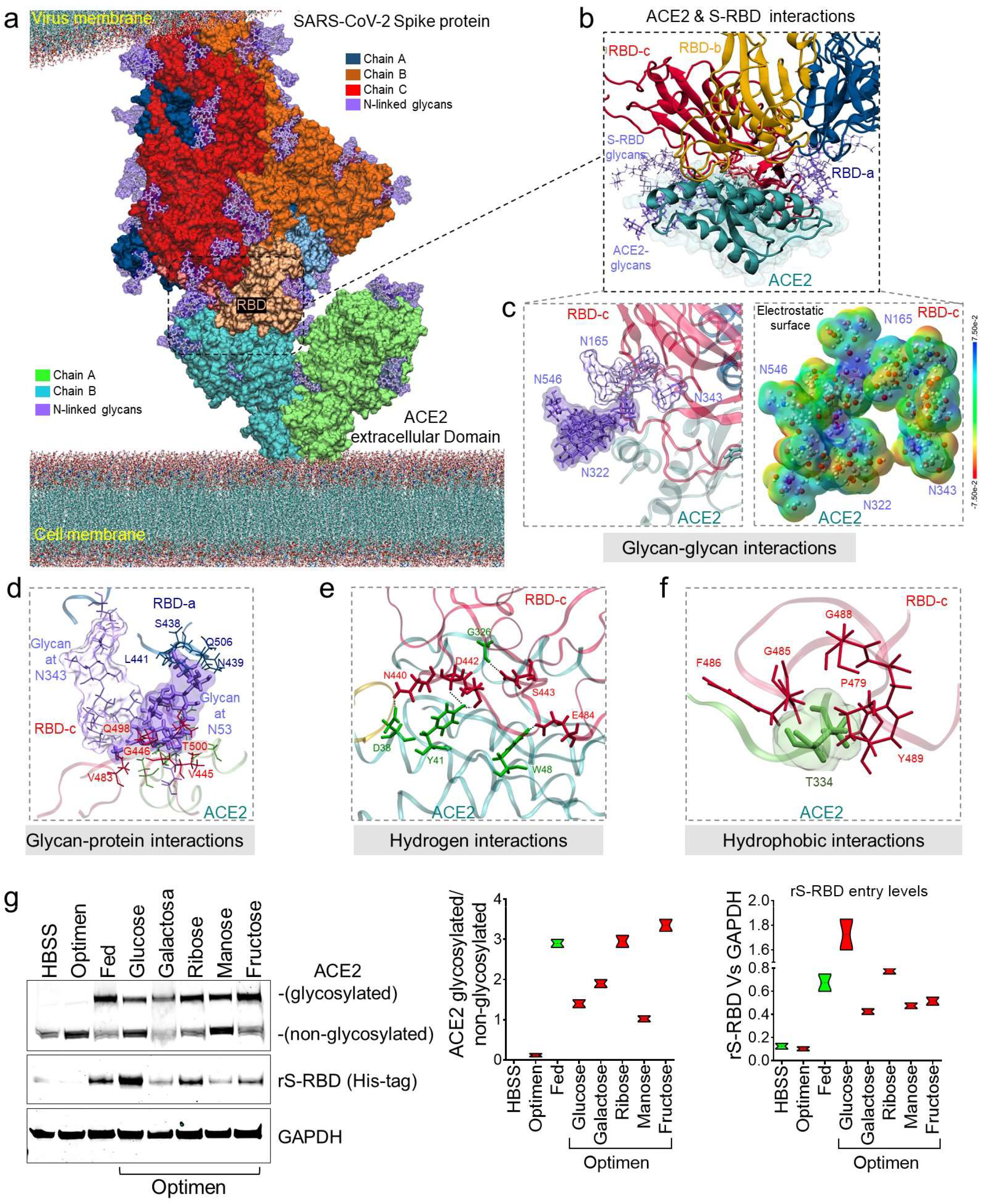
Molecular bases of glycosylated hACE2 and SARS-CoV-2 Spike protein complex. (a) 3D membrane surface representation of glycosylated ACE2 in complex with the SARS-CoV-2 Spike protein. (b) Close up of the interacting environment between ACE2 and the S-RBD trimer. (c) The left panel demonstrate glycan-glycan interactions between ACE2 (dark purple surface) and S-RBDc (light purple surface). The right panel shows that glycan-glycan contacts do not affect their molecular electrostatic potentials (MEPs) properties. The energy scale ranging from −0.075 μa (red) to 0.075 μa (blue). (d) ACE2 glycan at N53 forms glycan-protein contact with residues on the S-RBDa and S-RBDc proteins. (e) The ACE2 glycosylation induce the formation of hydrogen bonds that engages the helix α1 in the binding with multiple residues on the S-RBDc. (f) Hydrophobic interactions occur between ACE2 at T334 and multiple residues on the S-RBDc. (g) Immunoblot showing expression of glycosylated and non-glycosylated human ACE2 in HUVECs treated with difference saccharides. Glucose-treated cells exhibited the greatest internalization of the recombinant S-RBD. Quantification of protein levels of three replicate experiments is shown. Student’s T-test, 2 tails. Bar graphs are presented as mean with error bars (±SD).

### Estrogens bind to hACE2 and stimulate its stabilization and internalization

In an effort to explore the potential protective effects of female sex hormones against SARS-CoV-2 infection, we examined the impact of estradiol (17β-diol) and a dietary-derived phytoestrogen (S-equol) on hACE2 structure and protein expression by a combination of in silico modeling, in vitro, and in vivo analysis. Specifically, in light of the importance of glycan-glycan interactions that mediate virus-ACE2 interactions, we sought to analyze the effect of estrogens on key molecular viral and receptor binding sites. In agreement with Yan et al.^22^, we identified three important regions on the ACE2 surface that are utilized for SARS-CoV-2 binding. The environment of these regions is composed of a high density of glycans, including a helix α1 from residues I21 to T52, a helix α2 from residues V59 to M82, and one loop from residues K353 to G354 (suppl. Figure 3 and Figure 2a). We then homogenously solvated the glycosylated hACE2 structure with 26.6mM of 17β-diol or 26.5mM of S-equol followed by 100 ns of MDS. Remarkably we found that the 17β-diol molecules interact with residues at F28, Y41, Q76, T78, Q81, M82 and the S-equol molecules interacts with residues at Q24, K26, T27, F28, K31, E35, L39, N64, D67, K68, A71, F72, E75, Q76 and L79 (Figure 2b, supp. Table #). Both estrogen molecules energetically stabilized the α1 and α2 helices by physical interactions and thereby minimized the fluctuation of the ACE2 chains A and B (Figure 2c, supp 3#). Importantly, our calculation of free-energy landscape (FEL), demonstrated that the surface of chain B of ACE2 (S-RBD’s preferred interaction region) loses its interaction energy with the S-RBD protein from 10.2 kJ/mol to 8.58 kJ/mol (16%) for the 17β-diol system and to 9.18kJ/mol (10%) for the S-equol system (Figure 2d). In addition, binding of either estrogen molecules to the surrounding hydrophobic pocket of ACE2 at the residue T334 promotes a decrease in energy by ~12% which may have a negative impact on the attachment of the S-RBD protein to the receptor (suppl. Figure 4). We also observed estrogen-glycan interactions particularly at the glycan-protein interactions between the ACE2 (N53) and the S-RBD (N432) (Figure. 3a). Indeed, glycans are highly polar structures due to their high content of hydroxyl groups which make them suitable for attachment to the ACE2 protein (mostly negatively charged) or the SARS-CoV-2 S-protein (polarly charged). The density functional theory (DFT) calculation shows an important decrease of the glycan’s molecular electrostatic potential (MEP) due to the interactions with either estrogen molecules. Therefore, estrogen-glycan interactions could decrease the adhesive effect of glycans that enhance S-RBD and ACE2 receptor interactions (suppl. Figure 5 and Figure. 3b). These structural analyses suggest that estrogens could act as putative ACE2 ligands due to their ability to bind to highly energetic pockets at the top of the ACE2 surface protein which may increase its conformational equilibria and potentially boost its internalization to the cytoplasm. To support our in-silico analyses, we treated HUVECs with 17β-diol (3nM) or S-equol (10nM) overnight under normal physiologic conditions. Immunofluorescent staining demonstrated that estrogen-treated cells have less ACE2 membrane cellular localization (Figure 3c). Immunoblot analysis revealed that endogenous and dietary estrogens promote ACE2 internalization and degradation through the endocytosis process as assessed by LC3b^25^ and LAMP1^26^ protein activation in treated cells (Figure. 3d). To test the hypothesis that lower levels of estrogens are associated with increased levels of ACE2 protein in the respiratory tract, we administrated intratracheally either 17β-diol (0.3μM) or S-equol (1μM) to male mice. Histologic analysis of lung sections demonstrated that both forms of estrogens decrease ACE2 membrane expression levels in lung alveoli and also reduced the glycosylation of the ACE2 receptor (Figure 3e & 3f)

**Figure 2.**
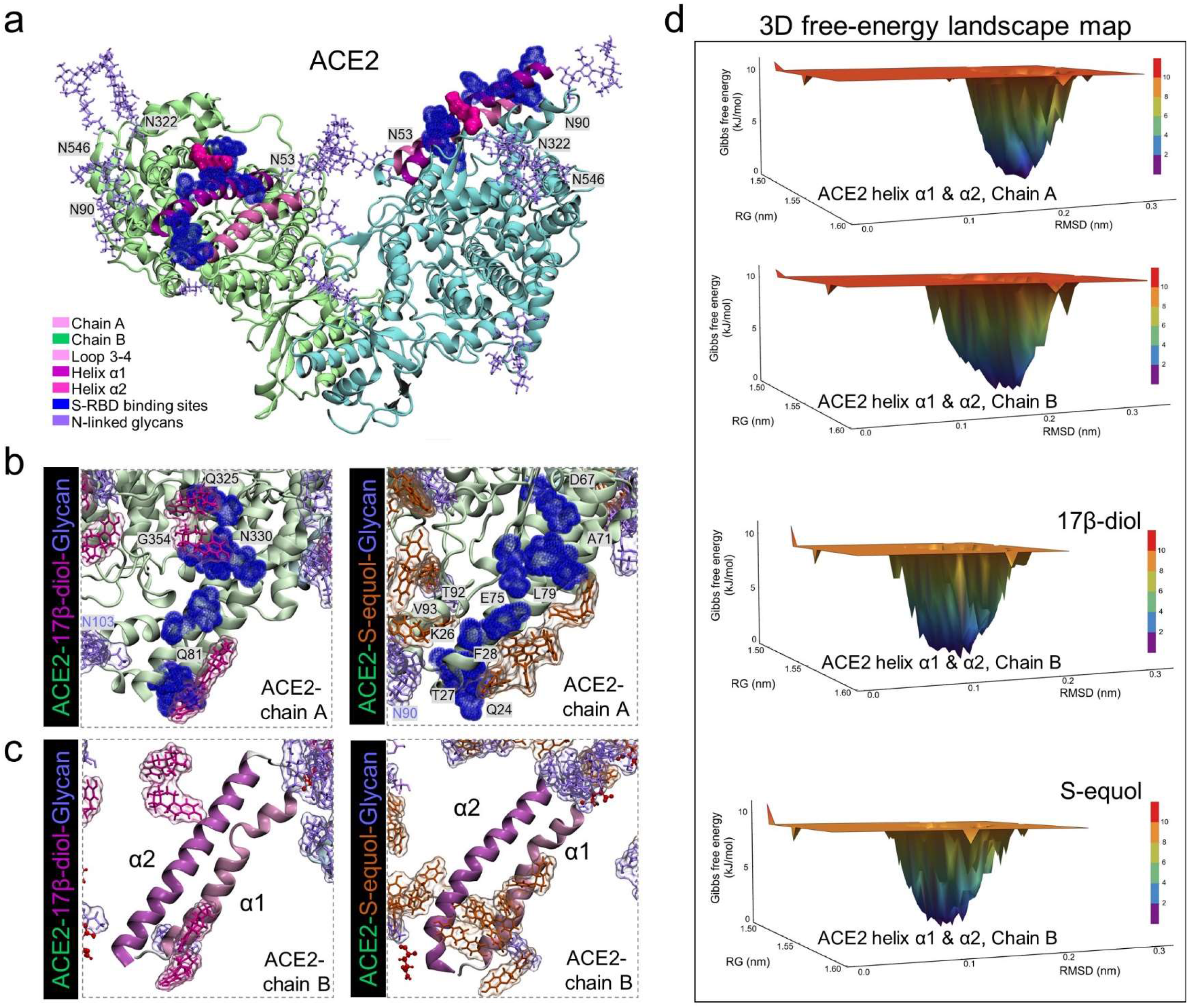
Estrogen effects on ACE2 structural energy. (a) 3D representation of the human ACE2 glycosylated residues and key regions used by the SARS-CoV-2 S protein to mediate entry into cells. S-RBD-binding sites are colored in dark blue and glycans in purple. (b) 3D molecular interactions between ACE2 and 17β-diol (magenta) or S-equol (organ) molecules obtained with 100 ns of molecular dynamics simulations (MDS). (c) A plain representation of solvated ACE2-helix α1 and α2 substructures by estrogen molecules. (d) FEL maps represent the conformational energy of helix α1 and α2 substructures with estrogen molecules during MDS (last 20 ns of MDS). Energy scale ranging from 12 kJ/mol (red) to 0 kJ/mol (blue).

**Figure 3.**
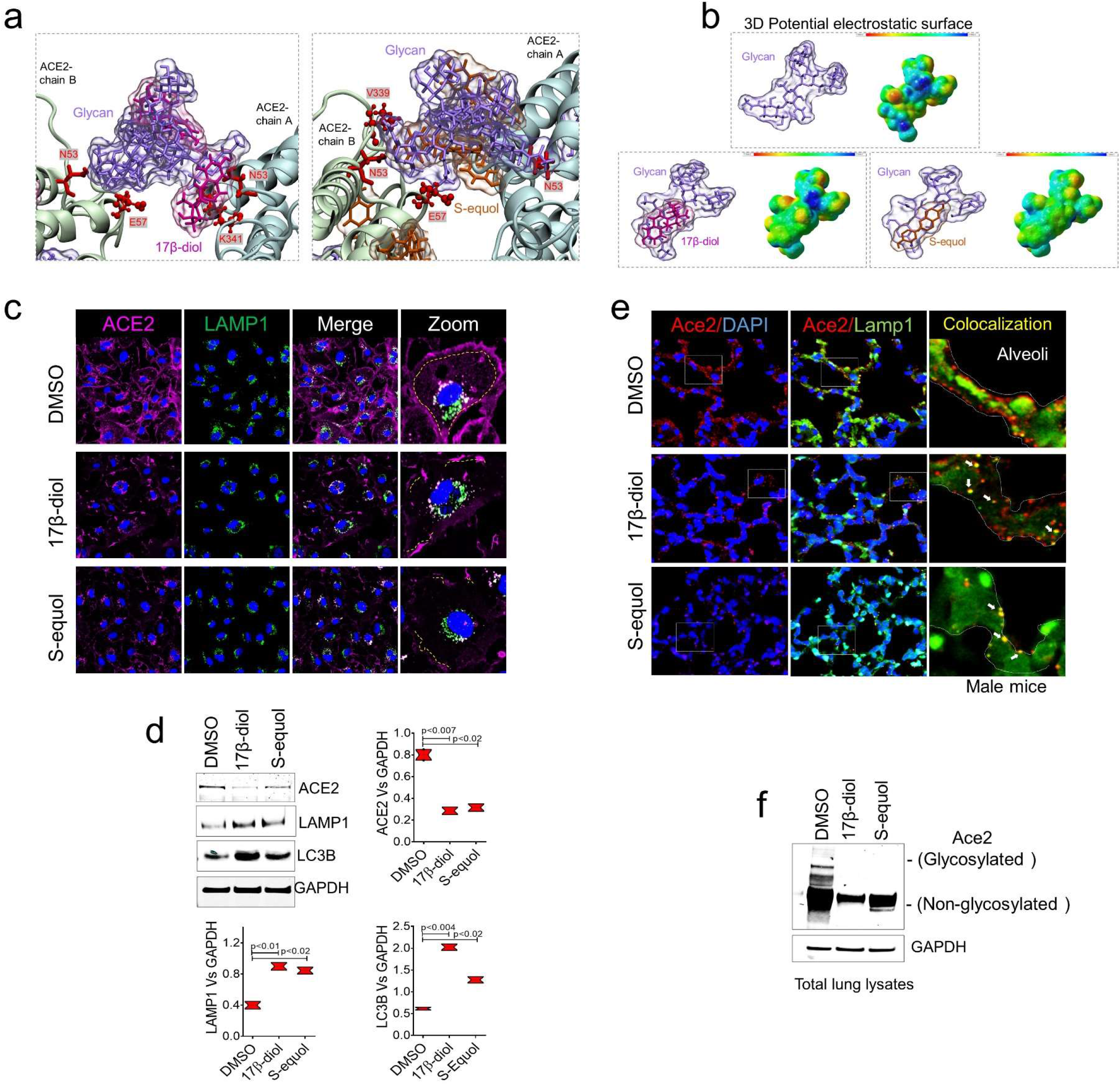
Estrogens bind to ACE2 glycans to promote its internalization. (a) Glycan-estrogen interactions stabilize ACE2 structure through high-energy contacts involving ACE2 glycan-residues at E57, N53, K341 and V339 (red color). (b) MEP maps show the electrostatic impact of estrogen molecules on the surface of ACE2 glycans. Energy scale ranging from −0.075 μa (red) to 0.075 μa (blue). (c) Immunofluorescence staining of human ACE2 (magenta) and the lysosome marker LAMP1 (green), shows loss of ACE2 membrane levels in HUVECs treated with 17β-diol or S-equol compared with control cells (DMSO). (d) Immunoblot of lysates isolated from HUVECs showing decreased levels of total ACE2 protein with estrogen treatment. Reduced ACE2 protein levels were associated with increased endocytosis activity as evidenced by immunoblot for LC3b and LAMP1. (e) Histologic analysis of mouse lungs after 48hrs of intratracheal installation with 17β-diol or S-equol shows loss of Ace2 expression (red) on the membrane of alveolar cells. Estrogen-treated lungs showed greater Ace2-Lamp1 colocalization (white arrows) indicating internalization of the receptor. (f) Immunoblot showing decreased levels of total and glycosylated Ace2 proteins in estrogen-treated lungs from male mice compared to control lungd. Quantification of protein levels of three replicate experiments is shown. Student’s T-test, 2 tails. Bar graphs are presented as mean with error bars (±SD).

### Estrogens interfere with SARS-CoV-2 receptor binding and block entry into the cell

To determine if the decline of conformational Gibbs free energy and gain in stabilization of ACE2 due to estrogen binding could affect the ability of the S protein to interact with the ACE2 receptor and thereby its entry into cells, we performed a refinement step of ACE2-free or ACE2-estrogen models with 100 ns of MDS followed by molecular docking with the SARS-CoV-2 S protein. From 241 structures obtained, 57 with top scores were chosen for further analysis (suppl. Table 4). The ACE2-17β-diol model promoted the shift of S-RBDs from the binding surface toward the lateral side of the ACE2 protein decreasing the number of contact residues. Notably S-RBDs completely lose contact with key ACE2-glycosylated residues at N53, N103, N432 and N690. We also observed that the contact between the S-RBD and the helix α1 and α2 of ACE2 moved toward the N-terminal of the helix and thus affected the ability to bind the receptor. In the same manner, the ACE2-S-equol model demonstrated that S-equol blocks the contact between the S-RBDs and the receptor’s surface, notably promoting novel interactions at the C-terminal of the helix α2 causing nonspecific contacts with the receptor at residues Q429-I436 and P590-N601. Interestingly, we found that the 17β-diol interacts with 66 residues on the surface of the receptor and notably forms a cluster on glycans at N546 (chain A) and N322 (chain B). On the other hand, the S-equol molecules tend to interact more widely accounting for a total of 145 interactions, including on 63 residues on the chain A and 82 residues on the chain B. (For better visualization, only the 5 top scored S-RDBs structures are shown in Figure 4a). The nonspecific binding by the S-RBDs could be explained by the susceptibility of ACE2 to interact with polar molecules and especially to electrophilic attacks. The fact that the 17β-diol or S-equol contain few polar groups but are deficient in negative charge renders them more susceptible to attack the surface of hACE2 thereby blocking S-RBD from binding correctly. In addition, we computed the binding score of these models using the atomic energy contact function and in agreement with our previous docking results observed that both estrogen molecules significantly reduced the atomic energy contact between virus and receptor. Remarkably, the 17β-diol reduced the atomic contact by 80% and the S-equol by 65% indicating that the entry of the virus may be affected by the presence of either estrogen molecules (Figure 4b). To validate our in-silico prediction, we pre-treated HUVECs with either estrogen molecules followed by incubation with 10μg of rS-RBD protein overnight. Importantly either low or high concentration of 17β-diol (low=1nM) & high=5nM) or S-equol (low=5nM & high=20nM) blocked more than 90% of the rS-RBD protein entry into the cell as assessed by immunofluorescence and colocalization with LAMP1, a lysosome marker (Figure 4c). In addition, immunoblot demonstrated a decrease of rS-RBD levels in the cytoplasm of HUVECs for both estrogen-based treatments (Figure 4d). Together these results suggest a potential molecular mechanism by which estrogens may provide protection against severe infection in COVID19 among women and individuals with phytoestrogen intake.

**Figure 4.**
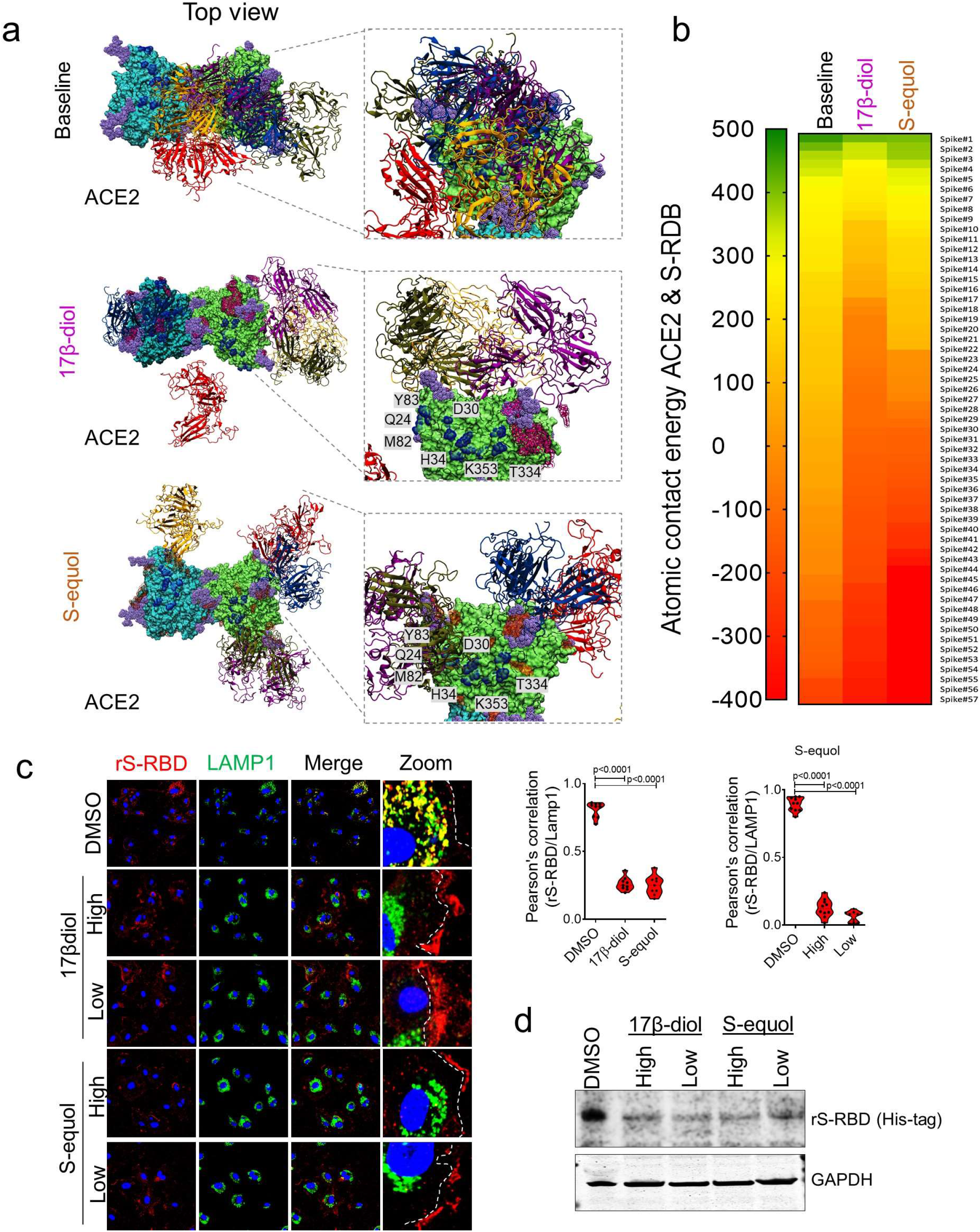
Estrogen impact on ACE2 and S-RBD interactions. (a) Top view of the 3D ACE2 surface interacting with 5 top scored S-RBDs (top 1-blue, 2-red, 3-orange, 4-purple and top 5-yellow). S-RBDs were scored based on shape complementarity principles. (b) Heatmap of atomic contact energy between ACE2 and 57 S-RBDs, shows spontaneous energy structures from most favorable (green) to less favorable S-RBD structures (red). Energy scale ranging from 500 Kcal/mol to −500 Kcal/mol. (c) Immunofluorescence analysis of S-RBD entry into HUVECs pre-treated with 17β-diol or S-equol followed by treatment with 10μg/mL of recombinant S-RBD (red) demonstrate that estrogen-treated cells had reduced entry of S-RBD into cells in conjunction with a reduction in ACE2 internalization as showed by colocalization with LAMP1 (green). (d) Immunoblot of isolated proteins from cultured HUVECs shows a 90% reduction of S-RBD entry into cells in estrogen-treated cells. Quantification of protein levels of three replicate experiments is shown. Student’s T-test, 2 tails. Bar graphs are presented as mean with error bars (±SD).

### Estrogens blocks SARS-CoV-2 infection of the respiratory tract *in vivo*

Next, we sought to test the ability of estrogens to block key interactions between ACE2 and the SARS-CoV-2 S protein and thereby infection of the respiratory tract. Male wild-type mice were treated with 17β-diol (0.3μM) or S-equol (1μM) via intratracheal instillation for 24hrs before tissue collection. ELISA-based binding assay showed a significant decrease of ACE2 affinity to SARS-CoV-2 S protein in lungs from mice treated with either estrogen molecules compared with the control group (Figure 5a). We then evaluated *in vivo* whether intratracheal estrogen treatment would reduce internalization of the S protein in male mice. We observed that intratracheal instillation of both estrogen molecules 24hrs before intratracheal instillation of rS-RBD (20μg, overnight treatment) increased signal for the rS-RBD on the surface of lung cells which likely results from reduced binding to Ace2 in estrogen-treated mice compared to the untreated group. In contrast control (DMSO-treated) lungs showed normal Ace2 membrane localization and cytoplasmic r-S-RBD signal indicating the unperturbed intake of the rS-RBD protein. The observed increase in extracellular rS-RBD in alveolar cells from lungs of estrogen-treated mice suggests estrogen-mediated reduction in internalization of the S-protein. Indeed, we observed that pretreatment with estrogen resulted in rS-RBD protein accumulation on the surface of the alveolar cells (Figure 5b) rather than being internalized into the cytoplasm which would thereby support viral replication and disease progression. Our data show that estrogens may interfere with SARS-Cov-2 infection in the respiratory tract through direct interaction with the Ace2 receptor *in vivo*.

**Figure 5.**
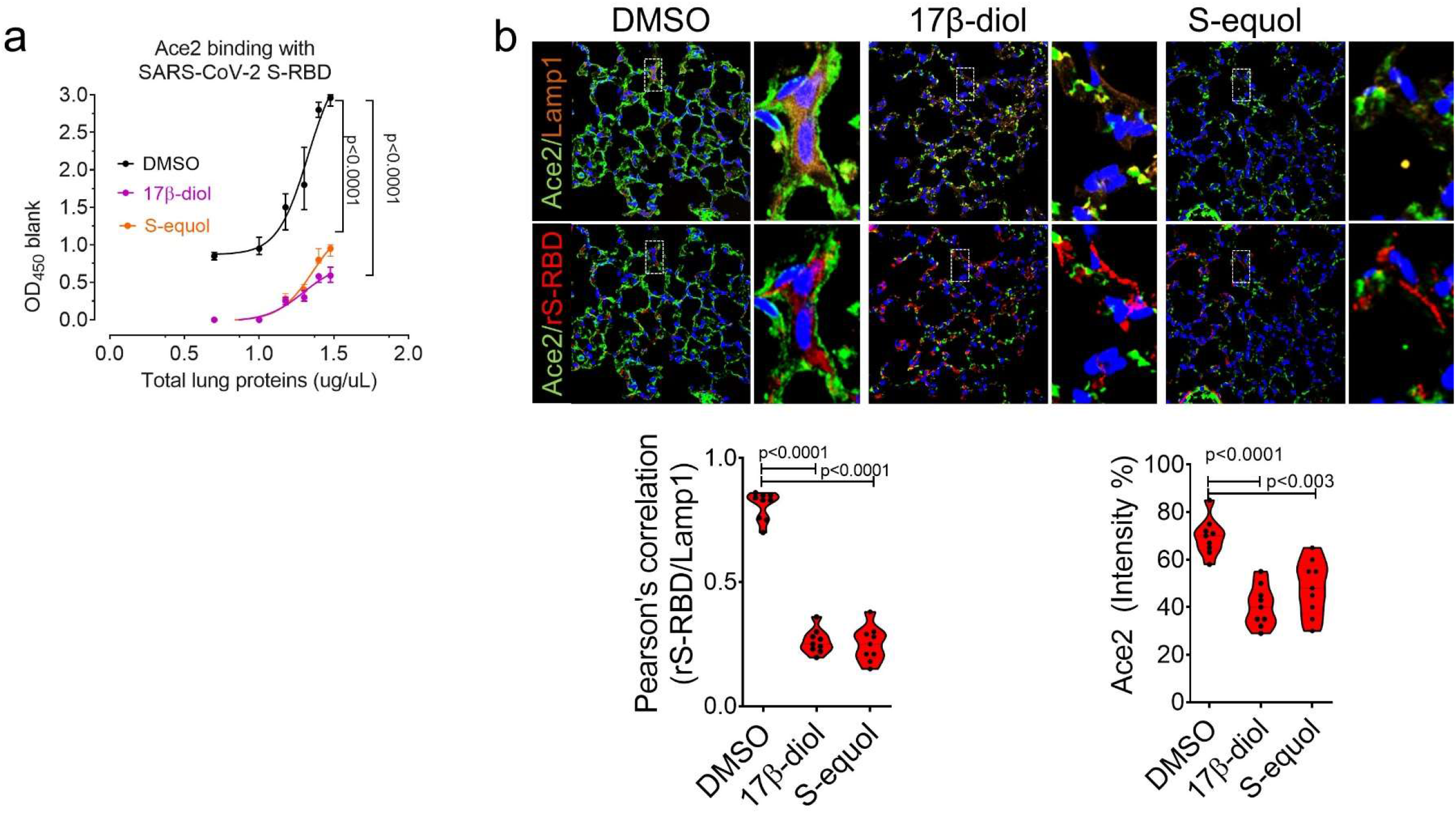
Estrogens blocks SARS-CoV-2 S protein uptake in the respiratory tract *in vivo*. (a) ELISA-based binding assay using lung protein lysates shows reduced SARS-CoV-2 S protein affinity for the Ace2 receptor after treatment with either 17β-diol or S-equol. (b) Immunofluorescence analysis of wild-type mouse lung treated with 17β-diol (0.3μM) or S-equol (1μM) compared with control lung (DMSO) demonstrates rS-RBD protein accumulation on the surface of the alveolar cells rather than being internalized intracellularly where viral replication may occur. Treatment with either estrogen also reduced Ace2 prtoein expression (quantification in lower right panel). Lamp1 (orange), Ace2 (green) and rS-RBD (red). Quantification levels of three replicate experiments is shown. Student’s T-test, 2 tails. Bar graphs are presented as mean with error bars (±SD).

## Discussion

Increased susceptibility and risk of adverse clinical outcomes among males affected by COVID-19 has been reported in multiple epidemiological studies^5–8^. Androgens can effectively upregulate viral target proteins that may increase viral entry and pathogenicity in patients following exposure to the SARS-CoV2 virus and sex-related hormones can modulate immune respose. A detailed understanding of the molecular and cellular mechanisms modulated by estrogen that contribute to viral pathogenicity is therefore critical to the development of new therapies to combat the COVID-19 pandemic. Beside the epidemiologic evidence suggesting that females are protected from severe infection, a recent study has demonstrated that the female reproductive tract, expresses very low levels of the ACE2 receptor and almost undetectable TMPRSS2, suggesting that the virus is unlikely to infect the female reproductive tract, where female sex hormones are produced^27,28^. In the current study, we utilized *in silico, in vitro*, and *in vivo* studies to characterize important glycosylation-mediated interactions between the SARS-CoV2 virus spike (S) protein and the human ACE2 receptor that can be modulated by endogenous or dietary estrogens in a manner that may be protective against the SARS-CoV2 entry into human cells.

Previous studies have highlighted the critical role of glycosylation in viral pathobiology, host immune system evasion, and infectivity in a range of human viral illnesses^29^. In many of these viruses, the viral envelope and secreted proteins are extensively glycosylated which is necessary for structural integrity and functionality of these proteins. Viral proteins may be glycosylated by the host cell as viruses are able to hijack cellular glycosylation processes. However little data exists on the impact of glycosylation of host proteins necessary for viral entry, such as ACE2, on viral infectivity. Using a novel molecular simulation approach, we demonstrated that ACE2 glycosylation augments binding of the viral S protein by supporting multiple types of interactions including glycan-glycan and glycan-protein interactions, thereby facilitating the stability and affinity of viral binding to the target host receptor. We extend these *in silico* observations by also demonstrating that entry of the S-RBD can be augmented *in vitro* by exposure of cultured HUVECs to a hyperglycemic environment that increases ACE2 glycosylation. These observations provide insights into the enhanced susceptibility of diabetic patients to severe infections and death in COVID-19^30–32^.

Based on these findings that ACE2 glycosylation enhances interaction with the viral S *protein in silico*, we explored whether the predominant endogenous form of estrogen, 17β-diol, may provide a protective effect as assessed using *in silico* modeling of viral S protein-ACE2 interaction and *in vitro* and *in vivo* models of viral infectivity. In addition, we used an identical approach to understand the potential protective mechanisms of dietary phytoestrogens on SARS-CoV2 infectivity observed in populations with low CFRs where consumption of these foods is high. We found that both endogenous and dietary estrogens compete with the S-RBD protein to bind specific sites on hACE2 that are used by the virus to bind the receptor. Indeed, estrogens were found to bind at almost all sites including ACE2 glycans causing a reduction of energy on the surface of the receptor rendering the receptor less susceptible to interact with other molecules via reduced cell surface expression including the viral S protein interactions. Our findings that estrogens interfere with S protein and ACE2 interactions *in silico* that is associated with reduced S protein uptake in an *in vitro* model of SARS-CoV-2 infectivity in cultured human endothelial cells are consistent with prior studies demonstrating that estrogens have antiviral properties against HIV, Ebola and hepatitis viruses^33^. Additionally, recent evidence indicates that decreased levels of estrogens in post-menopausal women are an independent risk factor for disease severity in female COVID-19 patients^34^. The findings of the current study thus represent novel findings in our understanding of the molecular mechanisms underlying reduced susceptibility to SARS-CoV-2 among females or individuals with depressed estrogen levels and in countries where dietary estrogens are high.

We then examined the ability of estrogen molecules to interfere with S protein uptake into pulmonary epithelial cells using an in vivo model of SARS-CoV2 infectivity. In agreement with our cellular experiments, lungs from mice treated with dietary or endogenous estrogens demonstrated a dramatic reduction in the uptake of S-RBD. In addition, we observed a remarkable reduction of ACE2 binding possibly due to the low protein levels of ACE2 in those lungs possibly in response to estrogen-mediated degradation. In conclusion we provide a molecular basis that helps elucidate the potential protective effect of estrogens in women infected by the SARS-CoV-2 virus which could inform the development of future therapeutic measures to protect against SARS-CoV-2 infection including the design of suitable blocking antibodies, estrogen-related treatments, and vaccine development.

## Acknowledgments

Dr. Lino Cardenas is supported by the MGH Physician-Scientist Development Award and the MGH Department of Medicine Pilot Translational Research Grant.

Dr. Malhotra is supported by a COVID-19 Fast Grant (fastgrants.org), NHLBI R01 HL142809, the American Heart Association grant 18TPA34230025, and the Wild Family Foundation.

Dr. Lindsay is supported by the Fredman Fellowship, the Toomey Fund for Aortic Dissection Research, the Hassenfeld Fellowship and NIH/NHLBI R01HL130113.

Dr. Davila-Del Carpio, Dr. Aguilar Pineda and Dr. Vera Lopez are supported by the COVID-19 Fast Grant (0371-VRINV-2020) from the Vicerrectorado de Investigación de la Universidad Católica de Santa María, Arequipa, Peru.

## Author contributions

Conceptualization, C.L.L.C. Methodology, C.L.L.C. Investigation, C.L.L.C., R.M., J.A.A.P., M.A., W.J., K.V.L. Resources, R.M., M.E.L., G.D.D.C., B.G.V. Data curation, C.L.L.C., J.A.A.P. Writing-Original Draft, C.L.L.C., M.A., R.M., M.E.L. Funding acquisition, C.L.L.C., G.D.D.C., R.M.

## Methods

### immunofluorescence microscopy

For immunefluorescence, HUVEC cells were cultured into 8-well Lab-TekTM II Chamber Slides (NuncTM) and were then treated with either 17β-diol at 3nM or S-equol at 10nM. Cells were rinsed twice with ice-cold PBS, fixed with 4% paraformaldehyde in PBS (PFA, Boston Bioproducts) for 10 min at rt, and were permeabilized with 0.1% Triton-X (Sigma–Aldrich) for 3 min. The slides were blocked with 10% Donkey-serum, and 0.3 M glycine in PBS-Tween 20 (0.1%) for 1 h at rt. Subsequently, the antibodies anti-ACE2 (1:100), S-RBD-His-tag (1:50), anti-LAMP1 (1:50) and anti-LC3b (1:50) were added and slides were incubated overnight at 4°C. The slides were then washed 3 times for 5 min each with PBS-T and were incubated with secondary antibodies at 1:400 dilution for 1hr at room temperature. Following immunostaining, slides were mounted with diamond mounting medium containing DAPI (Thermo Fisher). Slides were then visualized with the Leica TCS SP8 confocal microscopy station and the pictures were digitized with the Leica Application Suite X software.

### Protein Extraction and Immunoblot

HUVEC cells were rinsed twice with ice-cold PBS and proteins were extracted with M-PER for whole cell lysis, respectively (Thermo Fisher). These lysis buffers contained Halt protease, phosphatase inhibitors and EDTA (Thermo Fisher). The protein concentration was determined by the colorimetric bicinchoninic acid assay (BCA assay, Thermo Fisher). Equal amounts of total protein from cell lysates were separated by SDS-PAGE (25μg or 40μg for ACE2, LAMP1, LC3b and rS-RBD-His-tag, respectively). Proteins from the gel were then electro-transferred onto 0.45 μm nitrocellulose and 0.2 μm PVDF membranes. The membranes were then blocked for 1 h at room temperature, with either 5% non-fat powdered milk dissolved in TBS-T or 5% bovine serum albumin in TBS-T, for the nitrocellulose and PVDF membranes, respectively. Following blocking, membranes were incubated overnight at 4°C with the primary antibodies anti-ACE2 (1:1000), anti-LAMP1 (1:2000), anti-LC3b (1:1000) and anti-His-tag (1:1000). The Odyssey infrared western system was used to detect target proteins. Band intensity was quantified using ImageJ software.

### Animal treatment

All experiments involving mice were approved by the Partners Subcommittee on Research Animal Care. Personnel from the laboratory carried out all experimental protocols under strict guidelines to insure careful and consistent handling of the mice.

#### Mouse model of SARS-CoV-2 S protein entry

9 weeks old male C57BL/6 were purchased from The Jackson Laboratories, USA. To induce the recombinant S-RBD protein. Briefly, mice were anesthetized with sevoflurane inhalation (Abbott) and placed in dorsal recumbency. Transtracheal insertion of a 24-G animal feeding needle was used to instillate estrogen molecules, rS-RBD or vehicle (DMSO), in a volume of 80 μl. Mice were sacrificed 24 hrs after instillation of rS-RBS and lungs were removed for further analysis.

#### Histology

Lungs were then fixed in formalin (10%) for 24 hours before transfer to 70% ethanol for photography prior to paraffinization and sectioning (7 μM) paraffin embedding. Slides were produced for tissue staining) for quantitative analysis.

### In vitro treatment

#### Saccharides treatment

Hypoglycemic media was composed by HBSS buffer or Optiment media. Normal media contained complete endothelial cell growth media. For hyperglycemic media, Optiment was supplemented with D-glucose at 25mM, D-galactose at 50mM, D-ribose at 250μM, D-mannose at 300μM, or D-fructose at 20μM. HUVECs at 60%–70% confluence were supplemented with hypoglycemic, normal or hyperglycemic media 24 hours before incubation with 10μg of recombinant S-RBD-His-tag overnight.

#### Estrogen treatment

HUVECs at 60%–70% confluence were supplemented with opti-MEM 24 hours before treatment with complete growing media containing 17β-diol at a concentration of 3nM or S-equol at a concentration of 10nM for 24 hrs. Fresh media containing rS-RBD (10μg) was supplied the next day. Prior to cellular collection, cells were washed with sterile PBS, protein extraction were performed as described above.

### rS-RBD-ACE2 binding assay

500μg of total protein extracts from mouse lungs were cleaned up with IgA/IgG agarose beads for 1hr at 4C on a rotator followed by resuspension in assay diluent at 1x. Then 100μL of each lysate containing 0, 5, 10, 15, 20, 25 or 30μg of total protein were placed into corresponding well of a COVID19 S-protein microplate (Cat#: CoV-SACE2, Ray Biotech, inc.) for overnight incubation at 4C on a rotator. Then supernatant was removed, and wells were washed x 5 followed by incubation with 1x HRP-conjugated secondary antibody solution for 1hr at room temperature. Then 100μL of TMB one-step substrate reagent was added to each well for 30 min at room temperature. Before read 50μL of stop solution was added and microplate was read at 450nm.

### Statistics

Results are given as mean ± SD Student’s t test (2-tailed) was applied to determine the statistical significance of difference between control and treated groups (*P < 0.05, **P < 0.01 and ***P < 0.001). For all experiments, at least 3 experimental replicates were performed. Violin plot graphs show mean ± SD. Data were analyzed, and graphs were prepared with Prism 6.0 (GraphPad Software). P values of less than 0.05 were considered statistically significant.

### Model building and glycosylation process

The crystalline structures used in this work were PDB ID:6VXX^23^ for spike protein (SP, trimeric form) of virus SARS-CoV-2 and PDBID:6M17^22^ (of RBD/ACE2-B0AT complex) for ACE2 protein (dimeric form), both obtained in the RCSB Protein Data Bank. The missing residues for SP located on N and C terminal domains (M-18 to P26 and F1148 to H1262, respectively) were not considered in the molecular simulations. Therefore, each one SP chain was made up of 1121 residues (A27 to S1147). In our SP model, another disulfide bond was recognized between missing residues C499 and C507 and was considered in MD simulations. For ACE2, 29 residues on N-terminal domain were excluded (M-8 to T20) and on C-terminal domain only extramembrane residues were considered (to I21 to G732). ACE2 structure contain two zinc ions in peptidase domain which were considered in this work. The remaining missing residues of both proteins were added using SwissModel server^34^ (S-T1).

The ACE2 and SP structures are considered glycoproteins and the glycan-linked residues have already been reported^21–24^. The OPLSAA based DoGlycans software was used for building all models^35^ (S-T1). For SP, there are 22 N-glycosylation residues on its surface, but N17, N1158, N1173 and N1194 sites were excluded due to residues considered in our SP model. O-glycosylation sites was not included too. In ACE2, all N-glycosylation sites were considered. The glycosylation process was carried out using the glycan GlcNAc_2_Man_3_ model, a glycoside sequence composed of 2 N-acetyl glucosamines and 3 mannoses (S-F1b). This glycan type is the most common core sugar sequence on the N-glycans^36,37^

### Estrogen solvated systems

The systems were constructed with the average structure of glycosylated ACE2, obtained in last 50 ns of MDS trajectories. 17β-diol and S-equol structures were quantum optimized and their force fields were constructed using LigParGen server^38–40^. The previous simulation box of ACE2 was augmented 0.4 nm in all directions and with the protein centered in box, was solvated two times using *gmx solvate* module. The first solvation was made homogeneous way with 60 17β-diol and S-equol molecules (26.6 and 26.5 mM solutions, respectively). In second solvation, explicit water molecules were added to fill the simulation box. In the solvation process, we made sure that the estrogen molecules were not close to the protein at the start of the MD simulations.

### Simulations details

All quantum simulations were performed using density functional theory (DFT)^41^ at B3LYP/ TZVP level^42,43^. The self-consistent reaction field (SCRF) theory was used for describing the solvent effects on the molecules in water solutions. The calculations were performed in the electronic structure program Gaussian 16^44^ and results were visualized in GaussView v.6^45^. The molecular structures of 17β-diol and S-equol were optimized and it was ensured that they were at a global minimum through frequency analysis. These optimized structures were used in the molecular dynamic simulations. In MEPs analysis, single point calculations were carried out and total electron densities was mapped on molecular electrostatic potential surface.

To address the structural interactions, we performed molecular dynamics simulations using GROMACS (v.2019.4)^46^ with the OPLS/AA force field parameters^47^. The protein complexes were solvated with TIP4P explicit water model^48^. In addition to Na^+^ counterions used to neutralize the total charge in the simulation box, we used a 150 mM NaCl concentration to mimic physiological conditions. All molecular systems were built in a triclinic simulation box considering periodic boundary conditions (PBC) in all directions (*x*, *y* and *z*). Minimum distance of the surface atoms of proteins to the edge of periodic box was 1.5 nm for ACE2 receptor and SARS-CoV-19 spike protein, and 2.3 nm for ACE2-Estrogen solvated systems. The equations of motions were integrated with the Leap-Frog integrator^49^ using a time step of 1 fs. Temperature in the simulations was maintained at 309.65 K using modified Berendsen thermostat (V-rescale algorithm)^50^ with τ_*T*_ = 0.1 ps coupling constant with protein and water-ions coupled separately. Pressure was maintained at 1bar using the Parrinello-Rahman barostat^51^ with a compressibility of 4.5×10^−5^ bar^−1^ and a coupling constant of τ_*P*_ = 2.0 ps. All simulations were carried out with a short-range non-bonded cut-off of 1.1 nm and the particle mesh Ewald (PME) method^52^ was used for computing long-range electrostatic interactions with a tolerance of 1×10^5^ for contribution in real space. The Verlet neighbor searching cut-off scheme was applied with a neighbor-list update frequency of 10 steps (20 fs). Bonds involving hydrogen atoms are constrained using the Linear constraint solver (LINCS) algorithm^53^.

Simulations were first energy minimized using the *steepest descent* algorithm for a maximum of 100,000 steps. The equilibration was conducted by two steps. The equilibration was conducted by two steps. The first step, a 1 ns of dynamics in the NVT (isothermal-isochoric) ensemble and second step, was continued for another 2 ns in the NPT (isothermal-isobaric) ensemble. Production runs were performed in the NPT ensemble for 300 ns for ACE2 and SARS-CoV-19 spike protein and 150 ns for ACE2-Estrogen solvated systems.

### Structure and data analysis

The structural interactions were obtained carried out a rigid-rigid body docking analysis using PatchDock^54^ server in order to obtain the contact residues between S-RBD and ACE2 systems. The PatchDock algorithm discard all unacceptable complex and results are assorted by geometry shape complementarity score. In addition, PatchDock do calculate the effective atomic contact energies according to Zhang et al.^55^ The molecular docking was done take to ACE2 protein as receptor molecule and Spike protein as ligand molecule. Clustering RMSD value and complex type they were selected according to the recommended parameters for protein-protein interactions (4.0Ǻ and default mode). For ACE2-estrogen systems, the docking was performed in the presence of the estrogen molecules bound to the ACE2 structure, 40 17β-diol molecules and 47 S-equol molecules, respectively. From total results obtained in molecular docking, those structures that had steric impediments (intermembranal clashes) were discarded. The steric impediments were calculated based in SARS-CoV-2 virus size^56,57^, whose diameter varies about 60 to 140 nm and its spike protein is about 9 to 12 nm length (100 and 10 nm on average, respectively). Molecular interactions were analyzed with LigPlot software^58^ and the pdb files required was constructed with fortran based own computer programs and the statistical data results.

Statistical results, RMSD, RMSF, RG, SASA, hydrogen bonds, free energies, matches, structures, movies, b-factor maps, were obtained using Gromacs modules and their different tool options. The analysis of structure properties was performed using MD trajectories on the last 50 ns of each simulations and visualization of the MD simulations was created using Visual Molecular Dynamics (VMD) software^59^ and the graphs were plotted by the XMGrace software^60^.

Each molecular conformation during an MD simulation has an associated energy and this can be observed using FEL maps. These maps are usually represented by two variables related to atomic position and one energetic variable, typically the Gibbs free energy. This free energy can be estimated from probability distributions of the system with respect to the chosen variables that are then converted to a free-energy value by Bolzmann inverting multi-dimensional histograms. When represented in three dimensions, the FEL maps show the energy range of all possible conformations were obtained during a simulation. In this work, we considered two substructures of ACE2 protein for FEL map analysis, the Alpha1-2 region (I21 to Y83) and loops regions l2-3 and l3-4 (D303 to R357). The FEL maps were plotted using gmx sham module while the RMSD and radius of gyration were considered as atomic position variables respect to its average structure and figures were constructed using Wolfram Mathematica 12.1^61^.

---

**Suppl. Table 1.** Structural summary of the simulated proteins, Spike and ACE2. ^*a*^Total experimental residues reported. ^*b*^Per chain. ^*c*^ Total residues in computational simulations. ^*d*^ First and last residues proteins. The missing residues column shows the residues interval not located in the experiment.

| Spike and ACE2 glycoproteins data |  |  |  |  |
| --- | --- | --- | --- | --- |
| Protein | Total residues | Missing residues | N-linked Glycan (Asn) | Interaction sites |
| **Spike Trimer** | 3843<sup>a</sup> (1281<sup>b</sup>)<br>3363<sup>c</sup> (1121<sup>b</sup>)<br>(A27 to S1147)<sup>d</sup> | -18 to 26, 70 to 79,<br>144 to 185, 246 to<br>262, 445 to 488,<br>502, 621 to 640, 677<br>to 688, 828 to 853,<br>1148 to 1262 | 61, 74, 122, 149,<br>165, 234, 282, 331,<br>343, 603, 616, 657,<br>709, 717, 801, 1074,<br>1098, 1134 | R-B domains:<br>T333 - P527 |
| **hACE2 dimer** | 1628<sup>a</sup> (814<sup>b</sup>)<br>1424<sup>c</sup> (712<sup>b</sup>)<br>(I21 to G732)<sup>d</sup> | -8 to 20 , 769 to 805 | 53, 90, 103, 322,<br>432, 546, 690 | Q24, D30, H34,<br>Y41, Q42, M82,<br>Y83, Q325, E329,<br>K353, R357 |

**Suppl. Figure 1.**
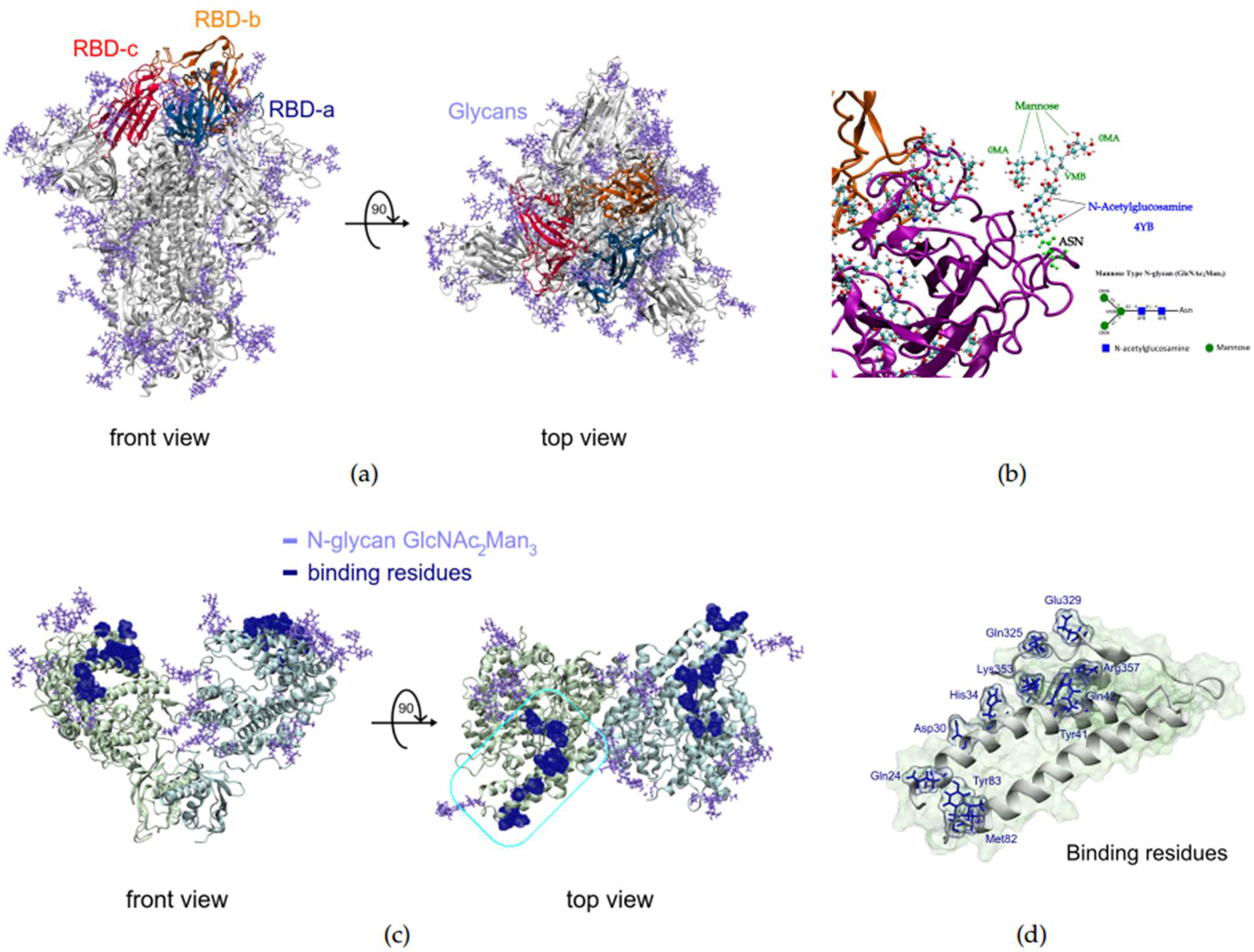
Structures building to experimental data and Doglycan software. (a) Spike protein, RBDs and glycans sites are shows.

**Suppl. Table 2.**

| Interaction | ACE2 Chain B | Spike Protein |  |
| --- | --- | --- | --- |
|  |  | Chain B | Chain C |
| Hydrogen bonds | Asp38, Tyr41, Trp48, Gly326 |  | Asn440, Asp442, Ser443, Asn450, Glu484 |
| Hydrophobic | Thr27, Lys 31, His34, Glu35, Ala36, Asp38, Leu39, Tyr41, Gln42, Leu45, Trp48, Asn49, Thr52, Asn61, Lys68, Phe72, Asp303, Ala304, Gln305, Thr324, Gln325, Gly326, Phe327, Trp328, Glu329, Asn330, Ser331, Met332, Leu333, Thr334, Asp335, Pro336, Gly337, Lys353, Asp355, Arg357, Ile379 | Ile472, Tyr473, Ala475, Gly476, Ser477, Thr478, Pro479, Cys480, Gly482, Val483, Glu484, Asn487 | Thr345, Arg346, Asn440, Asp442, Ser443, Lys444, Val445, Gly446, Asn448, Tyr449, Asn450, Tyr451, Pro479, Cys480, Asn481, Val483, Glu484, Gly485, Phe486, Cys488, Tyr489, Phe490 |

**Suppl. Figure 2.**
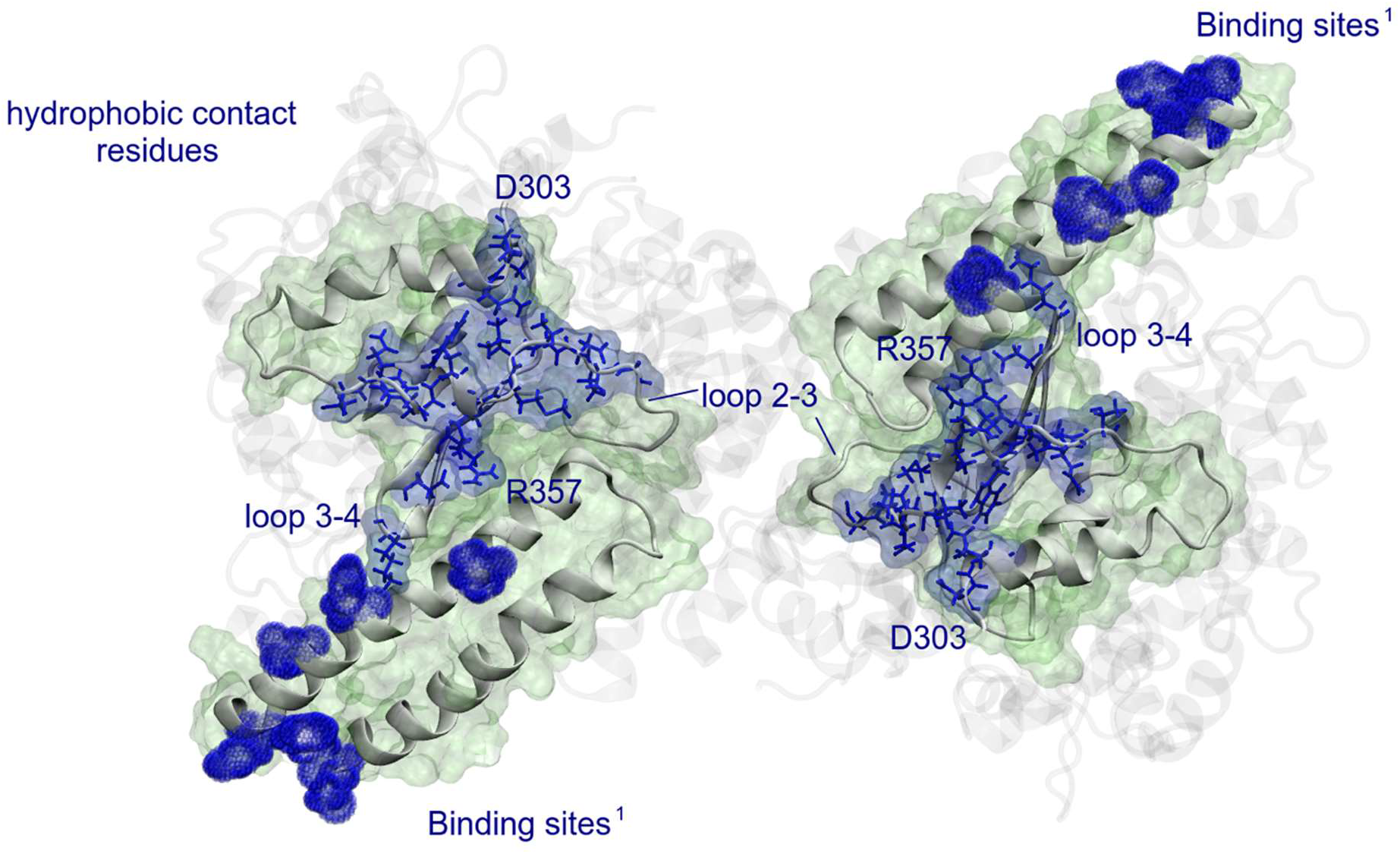

**Suppl. Figure 3.**
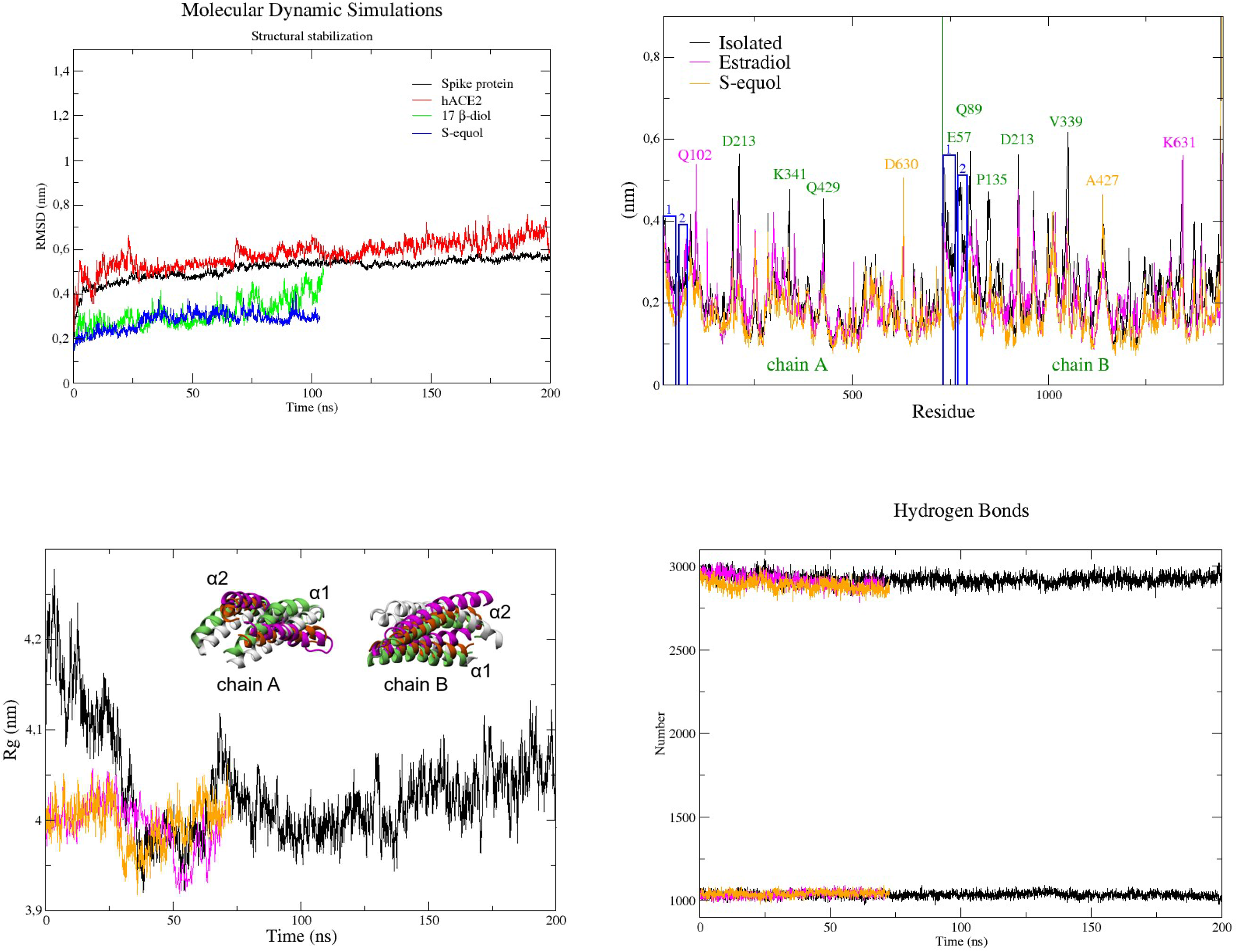

**Suppl. Figure 4a.**
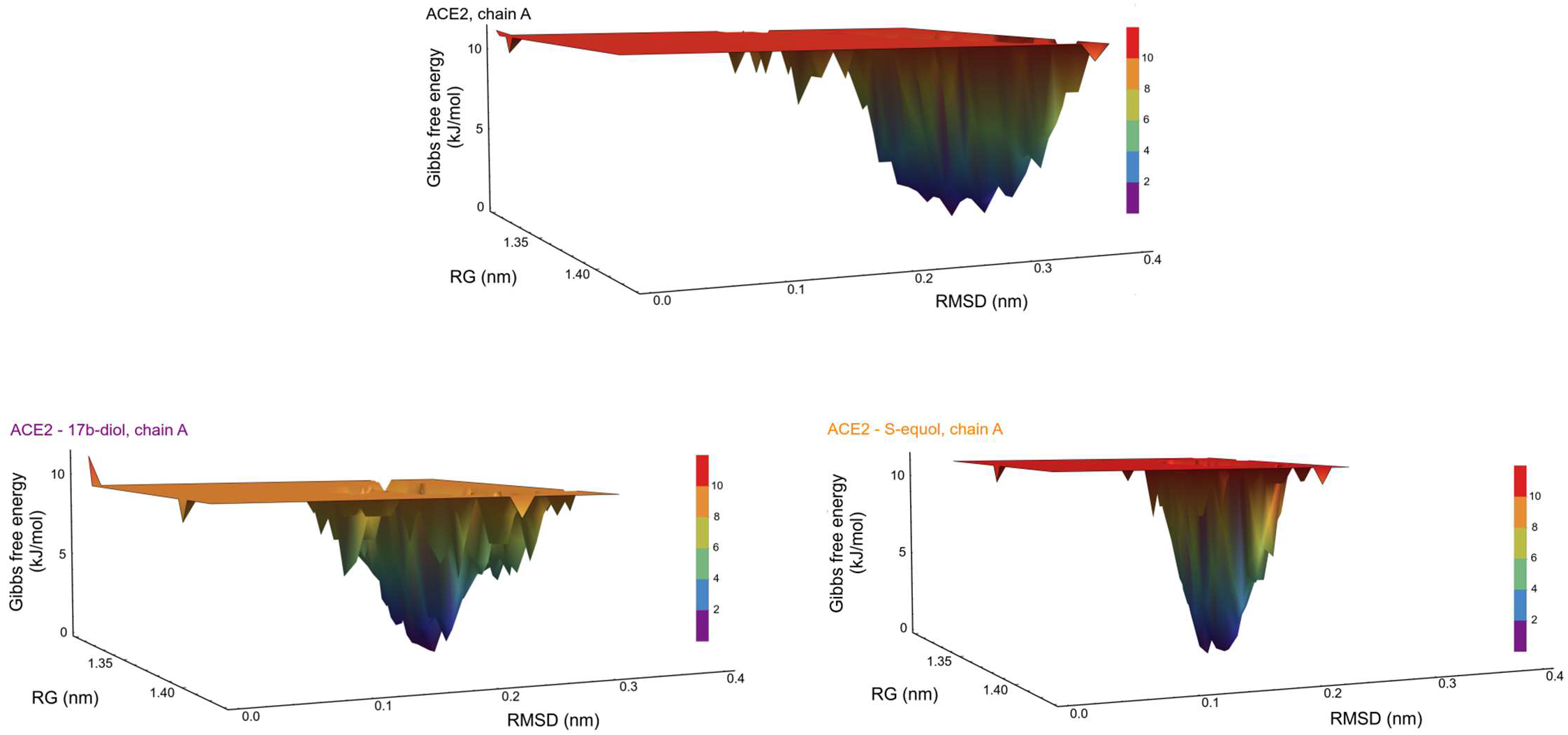
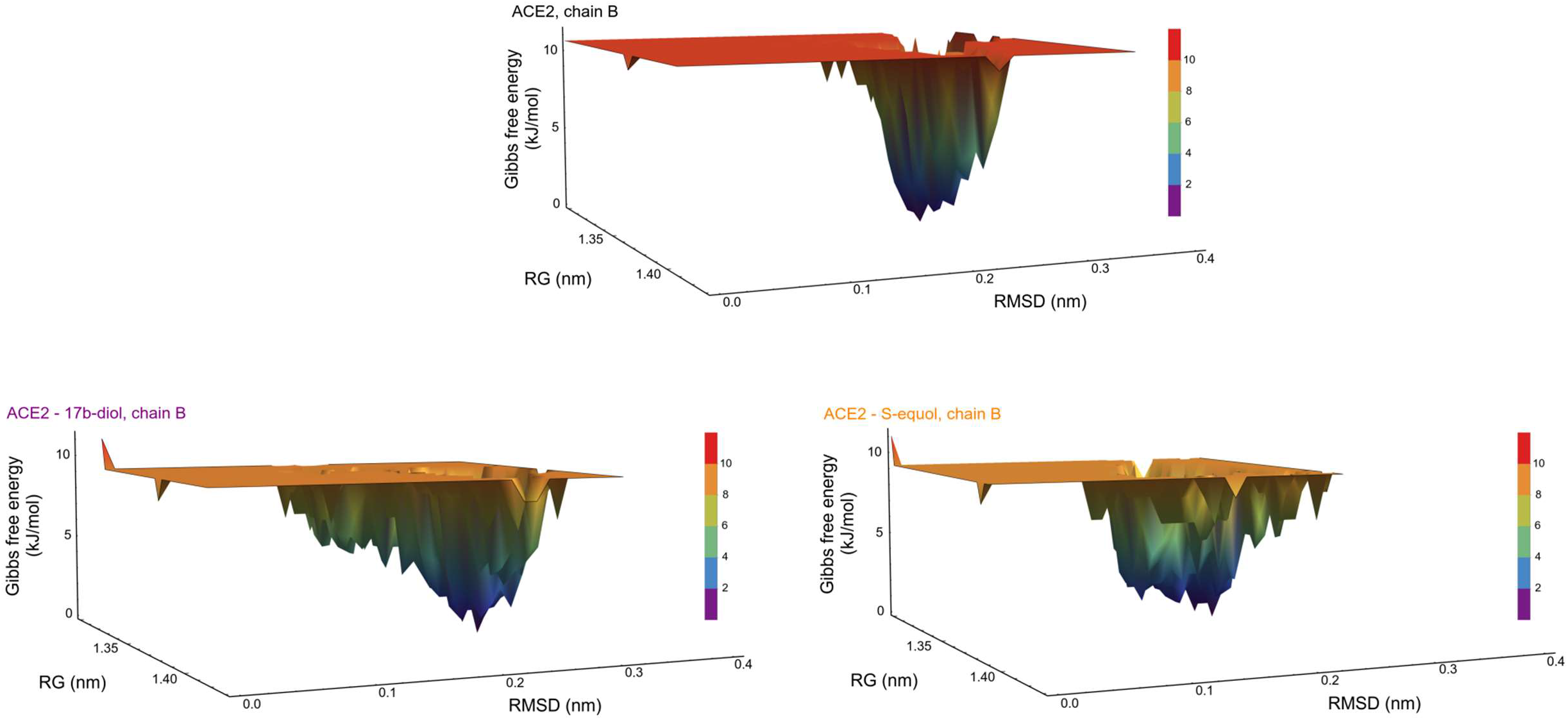

**Suppl Figure 5.**
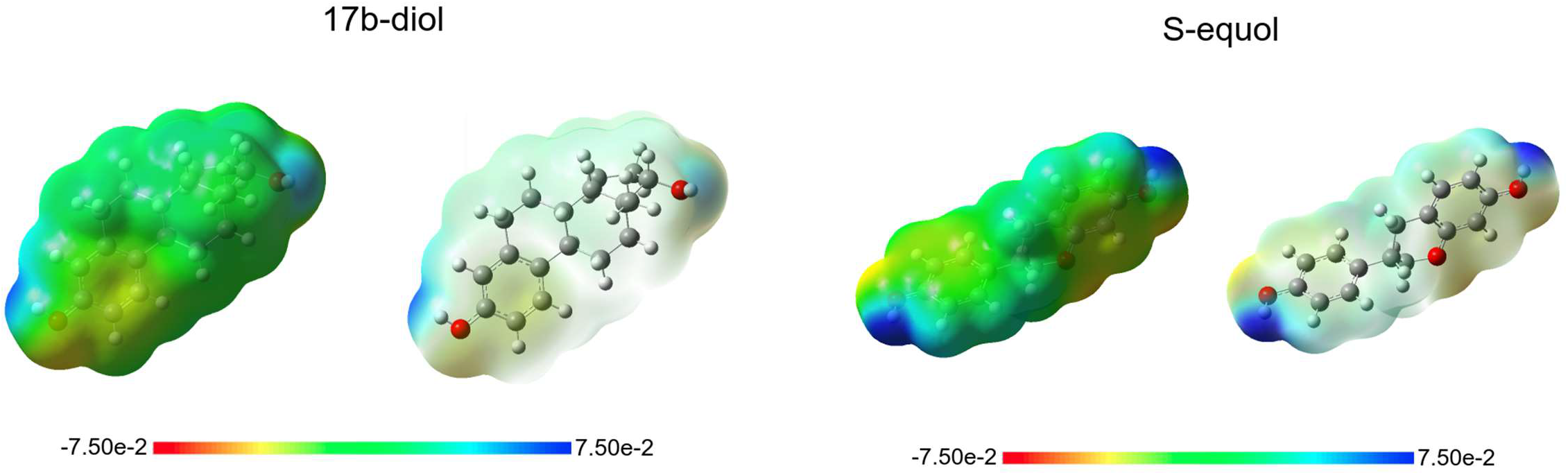

**Suppl. Table 4.**
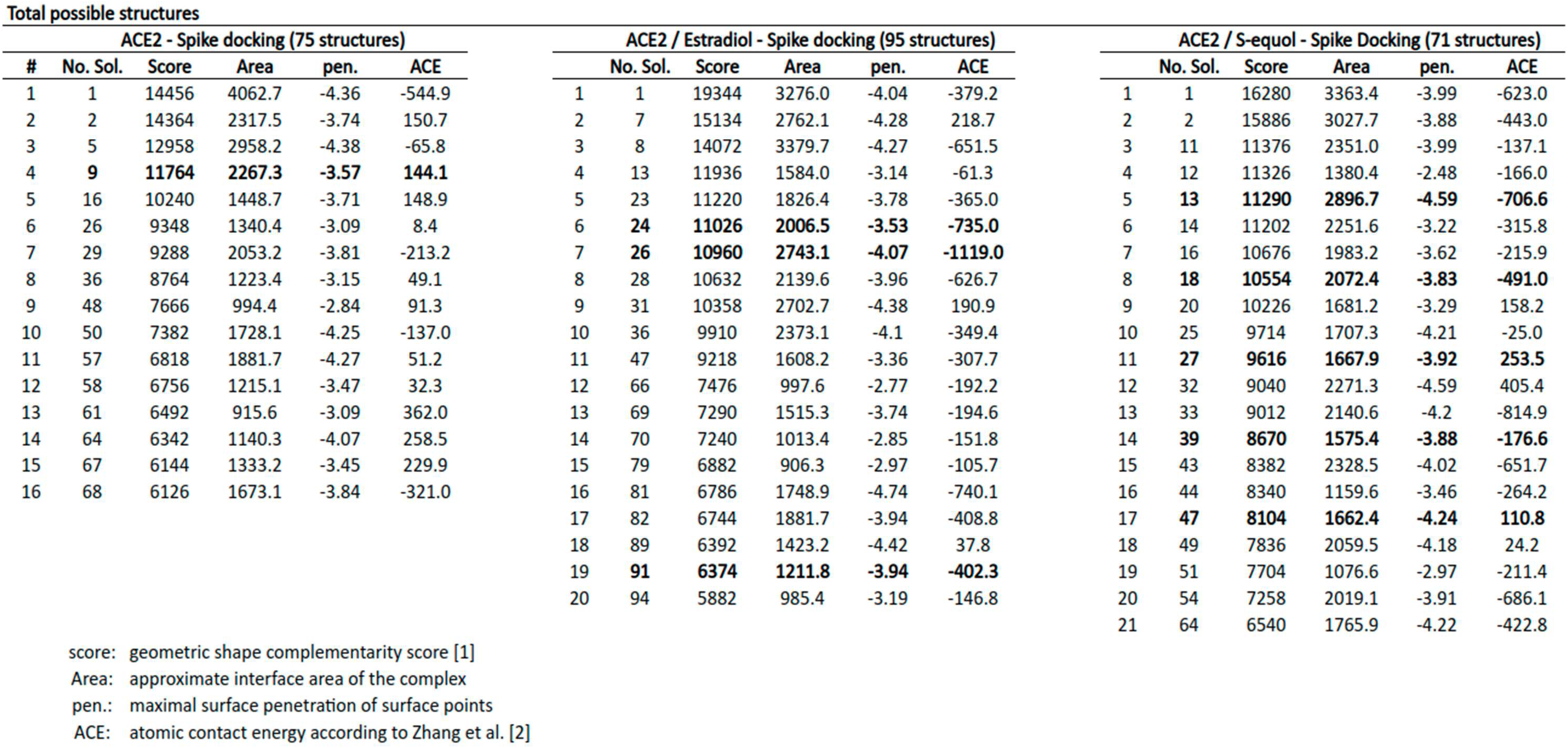

